# The Expression of HIV-1 Tat in *Lactococcus lactis*

**DOI:** 10.1101/2023.11.20.567837

**Authors:** Deepak Selvam, Anish D’silva, Yuvraj Gohil, Jayendra singh, Arun Panchapakesan, Luke Elizabeth Hanna, Udaykumar Ranga

**Affiliations:** Molecular Biology and Genetics Unit, Jawaharlal Nehru Centre For Advanced Scientific Research, Bengaluru, India; National Institute for Research in Tuberculosis, Chennai, India

**Author notes:** Corresponding author Email ID of the Corresponding author.

**Keywords:** HIV-1, Tat, P2 promoter, *Lactococcus lactis*, protein expression, endotoxin

## Abstract

Efficient expression of functional proteins in heterologous hosts has become the pivotal focus of modern biotechnology and biomedical research. To this end, multiple alternatives to *E. coli* are being explored for recombinant protein expression. *L. lactis*, being a gram-positive organism, circumvents the need for an endotoxin removal step during protein purification. We report here the optimisation of the expression of HIV-1 Tat, a notoriously difficult protein, in *Lactococcus lactis* system. We evaluated five different promoters in two different *Lactococcus lactis* strains and examined the effect of pH, glucose, and induction time on the yield and purity of Tat. Finally, the recombinant Tat was functionally competent in transactivating the HIV-1 promoter in HLM-1 reporter cells. Our work provides a scaffold for future work on the expression of toxic proteins in *Lactococcus lactis*.

## Introduction

Human Immunodeficiency Virus (HIV) is classified into two major types: HIV-1 and HIV-2. HIV-1, a more virulent and pathogenic virus, can be further categorized into several distinct subtypes based on genetic diversity[1] HIV-1 Tat, produced as an early protein during viral infection, is a basic protein of 14 to 16 KDa[2]. The primary function of Tat is to recruit P-TEFb to the viral promoter, the long terminal repeat (LTR), causing hyperphosphorylation of RNA pol II, leading to a substantial increase in the transcription rate[3]. Additionally, Tat modulates several cellular functions, including augmented cytokine and chemokine production[4], [5], cell signalling[6], splicing[7], DNA repair[8] and up-regulation of co-receptor expression[9].

A substantial amount of purified and functional Tat is required for in-depth characterization and in vitro investigations of this crucial viral protein. However, Tat can cause excessive cell death given its highly cytotoxic properties[9]. Although *E. coli* has been used as the preferred bacterial host for the expression of recombinant Tat[10], this system suffers from several technical limitations. The presence of several cysteine residues, typically 6 to 7, often causes protein misfolding, consequently leading to Tat sequestration in inclusion bodies. Further, the presence of bacterial Lipopolysaccharide (LPS), commonly referred to as endotoxin, in the purified protein preparation represents a major concern as it can independently upregulate cytokines and chemokines[11]. In our quest to circumvent these challenges, we explored expressing HIV-1 Tat in *Lactococcus lactis* (*L. lactis*), a gram-positive bacterium.

*L. lactis*, commonly employed in the dairy and cheese industry, has garnered significant attention in recent years in recombinant protein expression[12]. *L. lactis* thrives at a temperature of 30°C, proliferates with a rapid doubling time of 35 to 60 minutes, and exhibits both fermentative and respiratory metabolic pathways[13]. The ease of cultivation and cost-effectiveness of bacterial culture, combined with the availability of numerous genetic tools and vectors for recombinant protein expression, make *L. lactis* an attractive choice. Additionally, *L. lactis* offers the advantage of not forming inclusion bodies[12], [13]and the absence of endotoxin contamination. Furthermore, *L. lactis* can secrete proteins directly into the culture medium, simplifying the purification process. Douillard et al., 2011 successfully expressed a thioredoxin fusion protein at a concentration of 1.2 mg/l using the pTX8048 vector[14]. Bera et al., 2014 expressed an allergen protein under the control of the p170 promoter using the pMJ399 vector[15]. Notably, they also scaled up the culture from 1 l to 200 l. Martinez-Jaramillo et al., 2017 reported the expression of mCherry, GFP, and iRFP proteins in *L. lactis* using the pNZ8048 vector. They employed both pnisA and the p53 promoter for this purpose[16]. Finally, the moderate proteolytic activity of *L. lactis* ensures the integrity and proper folding of the recombinant proteins.

In the present study, we aimed to optimize the conditions for HIV-1 Tat expression in *L. lactis*, including selecting appropriate *L. lactis* strain, choosing the efficient bacterial promoter, optimizing the pH of the culture medium, modulating the glucose concentration, monitoring the harvest time.

## Materials and Methods

### The construction of the Tat-expression vector

We previously reported the cloning of full-length Tat from a primary clinical sample labelled BL43 into the pet21b+ plasmid (Siddappa et al., 2006). The Tat open reading frame (ORF) was amplified from pet21b+-BL43 Tat plasmid using the forward primer N4233 (5’GGAGACTTAGACTT**GGGCCC**TGCATATGGAGCCAGTAGATCCTAACCTA-3’) and the reverse primer N4234 (5’ATGGTATAC**AAGCTT**CTAGTGGTGGTGGTGGTGGTG-3’), and cloned into vector pNZ8148 using ApaI and HindIII restriction enzymes. The enzyme sites are bolded in the primer sequences. In vector pNZ8148, Tat expression was placed under the control of the nisA promoter. We generated a panel of Tat-expression vectors by substituting the nisA promoter with PepN, P2, and P11 promoters using the BamHI and ApaI sites. In the pAMJ399-Tat expression vector, the *tat* ORF was inserted downstream of the P170 promoter using BglII and PstI sites. MG1363 strain and plasmid pAMJ399 were obtained from Bioneer (Denmark), NZ9000 strain and plasmid pNZ8148 were obtained from NIZO (Netherlands).

### Electroporation of *L. lactis* and bacterial culture

*L. lactis* electrocompetent cells were prepared as described in MoBiTec GmbH (https://www.mobitec.com). Forty μl of *L. lactis* competent cells were dispensed into a pre-chilled 0.1 cm electroporation cuvette, and 1 μg of the appropriate plasmid was added to the cuvette. The electroporation conditions used were 1,000 V, 25 μF, and 200 Ω, resulting in a time constant of approximately 5 msec. Following the pulse, the cuvette was placed on ice for 5 minutes, and 1 ml of recovery medium (1X M17, 1% Glucose, 20 mM MgCl_2_, and 2 mM CaCl_2_) was added to the cells. The cells were then incubated at 30°C for 1 h and plated on an M17 agar plate supplemented with the appropriate antibiotic, chloramphenicol for pNZ8148 based vectors and Erythromycin for pAMJ399-based vectors. The bacterial plates were cultured at 30°C for 16 h.

A single bacterial colony was inoculated into 10 ml of M17 medium supplemented with 1% glucose and 10 μg/ml of the appropriate antibiotic. The cultures were incubated overnight at 30°C without shaking. A hundred microlitres of the primary culture were transferred to 10 ml of secondary culture. At an optical density (OD) of 0.4, the bacterial culture medium was supplemented with 1 ng/ml of nisin to induce protein expression from nisin-inducible vectors. After 4 h of induction, cells were harvested to assess Tat expression. Bacterial cultures harbouring the constitutive promoters P2, PepN, P11, and P170 were harvested 4 h after inoculation of the secondary culture. Cell pellets were resuspended in a 100 μl of solution containing 10 mM of Tris, 1 mM of EDTA, and 10 mg/ml Lysozyme, and the vials were incubated at 37°C for an hour. A hundred microliters of 20% SDS solution were then added to each vial and thoroughly mixed.

### Evaluation of Tat expression by Western blot

Bacterial cell lysate, a quantity equivalent to 0.4 OD of the original bacterial culture, was resolved on a 15% SDS-PAGE gel. The protein bands were transferred to a nitrocellulose membrane using a Bio-Rad wet transfer apparatus. The membrane was blocked with 5% skimmed milk for one hour at room temperature with gentle agitation and washed three times with 0.05% Tween-20 in 1X PBS. The blot was probed with an anti-His-tag antibody conjugated to horseradish peroxidase (HRP) (Abcam, Cat No. ab1178). Alternatively, the membrane was first incubated with an anti-Tat antibody (#4138, NIH AIDS Research and Reference Reagent Program) for one hour at room temperature, followed by probing with an anti-mouse antibody conjugated to HRP (Calbiochem, Cat. No. 401253) for one h and washed three times with 0.05% Tween20 in 1X PBS. The blot was developed using WesternBright™ ECL (Advansta Cat. No. K-12045-D20) and imaged using an iBright 1000 imaging system.

Alternatively, Tat expression from P2-Tat and P170-Tat vectors from *L. lactis*, strains NZ9000 and MG1363, was evaluated using a dot-blot strategy and a commercial antibody targeting the His-tag associated with Tat. Multiple colonies were analysed for Tat expression (data not shown), and the colonies showing high protein expression were further advanced to a secondary culture. After an induction period of eight hours, the bacterial cells were harvested and used for protein purification.

### Tat purification

The high-yield cultures identified in the dot-blot analysis were transferred to an eight-litre culture containing 10 μg/ml of chloramphenicol and incubated at 30°C for four hours or until an optical density (OD) of 0.4. The cells were harvested by high-speed centrifugation, resuspended in 50 ml of lysis buffer (50 mM NaH_2_PO_4_, 300 mM NaCl, 1 mM DTT, 2 mM PMSF, 0.4 mM EDTA, 5 mg/ml of Lysozyme, 0.1% IGEPAL, and 20 mM Imidazole), and incubated at 37°C for one h to facilitate cell wall lysis. Subsequently, the cells were sonicated for 5-second on and 5-second off cycles, for a total of 10 minutes, using a Branson Sonifier 450. The sonication cycle was repeated for two more rounds to ensure cell lysis.

Following sonication, the bacterial cell lysate was centrifuged at 24,000 rpm for 20 minutes at 4°C, to remove the cell debris. Five hundred microliters of pre-incubated Ni-NTA resin were added to the clear cell lysate and incubated at 4°C for four hours with mild agitation at 8 rpm. The resin was loaded to a column and washed with 50 ml of the wash buffer (50 mM NaH_2_PO_4_, 300 mM NaCl, 1 mM DTT, 2 mM PMSF, 0.4 mM EDTA, 0.1% IGEPAL, and 50 mM Imidazole). Tat was eluted in five distinct fractions using 500 μl of elution buffer (50 mM NaH_2_PO_4_, 300 mM NaCl, 1mM DTT, 2 mM PMSF, 0.4 mM EDTA, 0.1% IGEPAL, 250 mM Imidazole) each time. The eluted fractions were analysed using SDS-PAGE, and the fractions containing maximum yield were pooled and dialysed (1XPBS, 1mM DTT, 2 mM PMSF, 10% Glycerol). Fifty-microliter aliquots of dialysed Tat were stored at -80°C.

### Assessing the impact of pH and Glucose concentration on Bacterial Growth

Ten individual colonies of MG1363 strain harbouring the P2-Tat vector were inoculated in 10 ml of culture medium. Tat expression in each culture was confirmed using dot-blot analysis. The culture showing the highest Tat expression was expanded by transferring a 5 ml culture to 500 ml of M17 Medium (Hi-media Cat. No. M1907-500G). The pH of the culture was measured at two-hour intervals. Four conditions of culture were maintained. In one set, the pH was maintained at 7.0 by adding 5N NaOH and pulsing with 6.25 ml of 20% glucose to 0.25% final concentration every six hours. In the second set, the pH was adjusted to 7.0 using 5N NaOH but without glucose supplementation. The other two culture sets were analogous to the above with respect to glucose supplementation but without pH adjustment. Samples were harvested every two hours for Western blot analysis to assess levels of protein expression. In some experiments, we evaluated varying glucose concentrations (0, 0.25, 0.5, 1 and 2%).

### Tat integrity analysis by Mass Spectrometry

One microgram of Tat protein was dialyzed using ultrapure water to determine the intact mass of purified Tat protein. This purified protein was then analyzed using LC-HRMS (Q Exactive HF, Thermo Fisher Scientific). For the LC-HRMS analysis, 200 nanograms of the Tat protein were injected into a UHPLC system (Dionex Ultimate 3000) with an autosampler, and the separation was performed on a C8 column (Hypersil Gold, 100 X 2.1mm, particle size 5 microns, Thermo Scientific, USA). The mobile phases used were 0.1% (v/v) formic acid in water (Mobile Phase A) and 0.1% formic acid in acetonitrile (Mobile Phase B). The flow rate was set at 300 μl/min, and the following gradient was applied: 0-1 minutes, 10% B solvent; 2-8 minutes, 80% B solvent; 9-11 minutes, 95% B solvent; 12-15 minutes, 10% B solvent. The column temperature was maintained at 60°C through the experiment. The MS instrument was equipped with a Heated Electrospray Ionization (HESI-II) probe as the ion source. Parameters for the ion source included a spray voltage of +3.8 kV, a sheath gas flow rate of 20, an auxiliary gas flow rate of 10, an S-lens RF value of 80, a probe temperature of 200°C, and a capillary temperature of 320°C. The MS method involved full positive polarity MS scans at a resolution setting of 120,000, covering a mass range of 500-2000 m/z. The maximum analysis run time was set at 15 minutes.

### Functional validation of LTR transactivation by recombinant Tat

Fifty thousand HLM-1 cells were seeded into each well of a 24-well culture plate. The cells were incubated with 1 μg/ml of recombinant Tat protein for 4 hours in a serum-free MEM medium (HI-Media Cat. No. AL047S-500ML). The serum-free medium was replaced with 5% MEM, the plates were incubated for 48 hours, and the culture supernatant was collected for analysis. Empigen was added to each vial to a final concentration of 0.1%, and the samples were incubated for 20 minutes at 56°C to inactivate the virus and release the p24 antigen. The amount of p24 present in each sample was determined using a 4^th^ Generation Microlisa HIV Ag & Ab commercial kit following the instructions provided by the manufacturer (J. Mithra Cat. No. IR232096, India). In an alternative cell model, fifty thousand TZM-bl cells were seeded into each well of a 24-well culture plate for 24 hours. Prior to transfection, the cells were washed with 1X PBS to remove residual serum proteins. Cells were transfected with 1 μg/ml of recombinant Tat using a commercial transfection reagent Pro-ject Protein Transfection Reagent Kit (Thermo Scientific Cat. No. 89850) following the manufacturer’s instructions. The cells were lysed with 1X passive lysis buffer (Promega Cat. No. E1941) and 10 μl of cell lysate, and 10 μl of Bright-Glo Luciferase (Promega Cat. No. E2620) substrate added, and the Luciferase value measured using 2450 Microplate counter from PerkinElmer, USA.

## Results

### 3.1 Identifying an optimal bacterial promoter for Tat expression

The nisin-induced gene expression system has been a popular model for expressing foreign proteins in gram-positive bacteria. We, therefore, cloned the HIV-1 *tat* ORF into the pNZ8148 vector[17] downstream of the nisA promoter using ApaI and HindIII sites. When the nisA promoter was induced with 1 to 5 ng/ml of Nisin (Data not shown), we observed a very small amount of Tat protein (∼2 μg/l) expressed, although this promoter is known to be one of the strongest promoters in *L. lactis*[18]. Therefore, we evaluated several other inducible (P170[19]) or constitutive (P2[20], P11[21], and pepN[22]) bacterial promoters for efficient expression of Tat. The original nisA promoter in the pNZ8148-BL43-Tat vector was replaced with PCR-amplified P2, P11, or PepN promoters to generate a panel of Tat-expression bacterial vectors (Fig. 1A, left panel). We also cloned the *tat* ORF into the pAMJ399[23] vector to place Tat expression under the inducible P170 promoter using the BglII and PstI sites (Fig. 1A, right panel). Bacterial cultures were harvested four hours after the secondary inoculation, the cells were centrifuged, treated with lysozyme for one hour at 37°C, and subjected to sonication for cell lysis. Cell lysates equivalent to 0.4 OD were resolved on a 15% SDS-PAGE gel, and protein fractions were transferred to a 0.2 μm nitrocellulose membrane and subjected to the Western blot analysis using an anti-His-tag antibody (Fig. 1B). We observed the highest protein expression from the P2 promoter followed by the P170 promoter. A relatively low-level Tat expression was evident from the P11 promoter, whereas no protein expression was seen from the nisA and PepN promoters.

**Figure 1:**
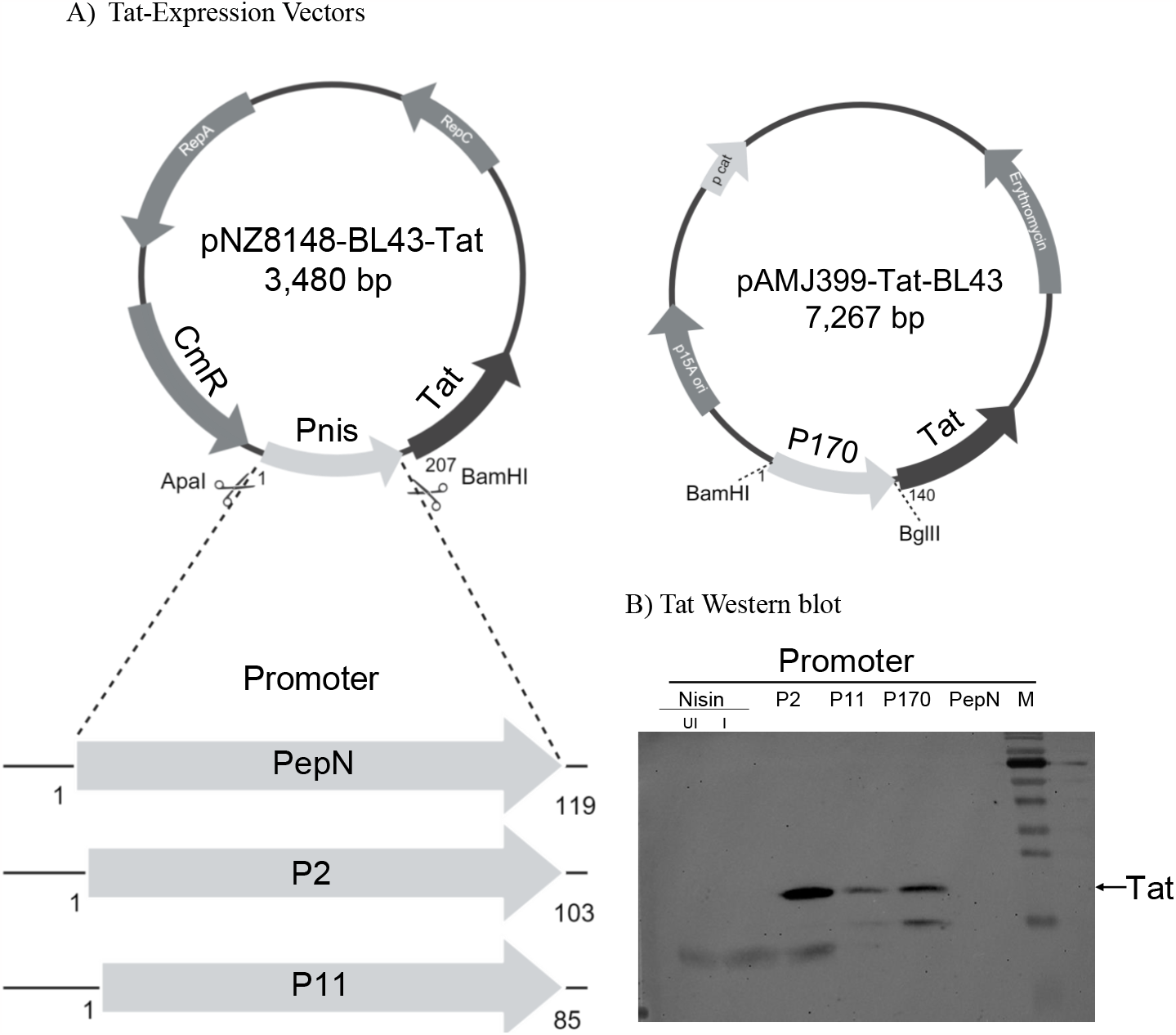
HIV-1 Tat expression in *L. lactis*. **(A)** Schematic representation of HIV-1 BL43 Tat expression vectors. The *tat* ORF of HIV-1 subtype C clone BL43 was PCR-amplified and cloned into the pNZ8148 plasmid backbone under the control of the pnisA promoter. The pnisA promoter was subsequently replaced with the P2, PepN, or P11 promoter using the specified restriction enzyme sites. The *tat* ORF was also cloned into the pAMJ399 vector downstream of the P170 promoter. **(B)** Tat expression in the NZ9000 strain of *L. lactis* from different bacterial promoters. Bacterial cultures of 10 ml were harvested four hours following the secondary inoculation. The bacterial cells were centrifuged, treated with lysozyme for one hour at 37°C, and subjected to sonication to lyse the cells. Cell lysates equivalent to 0.4 OD were separated using a 15% SDS-PAGE gel, and the protein was transferred to a 0.2 μm nitrocellulose membrane and subjected to the Western blot analysis using an anti-His-tag antibody. The Nisin promoter was induced with 5 ng/ml of Nisin, the P170 promoter was upregulated in response to lactic acid accumulation in the growth medium, while the P2, PepN, and P11 promoters are constitutive promoters.

### 3.2 The effect of Bacterial strain variation on Tat expression

Since the P2 and P170 promoters mediated efficient Tat expression in the NZ9000 strain of *L. lactis*, we compared protein expression from these two promoters in a second bacterial strain, MG1363. Tat was expressed in eight-litre cultures of the two bacterial strains under the two bacterial promoters, cells were harvested, recombinant Tat protein was purified from the cell lysates using Ni-NTA chromatography, and the protein expression levels were compared in SDS-PAGE. The data showed that Tat expression in strain MG1363, compared to NZ9000, offered two technical advantages a higher-level protein expression and low-level protein degradation (Fig. 2). Of note, adding a protease inhibitor cocktail to the culture medium did not prevent Tat degradation. In summary, we observed relatively higher and superior levels of Tat expression in the MG1363 strain of *L. lactis*, especially from the P2 promoter. We obtained 0.5 mg/l of protein in the eluted fractions, and there was no endotoxin contamination (data not shown). We used this expression strategy in subsequent experiments.

#### A) SDS-PAGE Analysis of the Purified Tat

**Figure 2:**
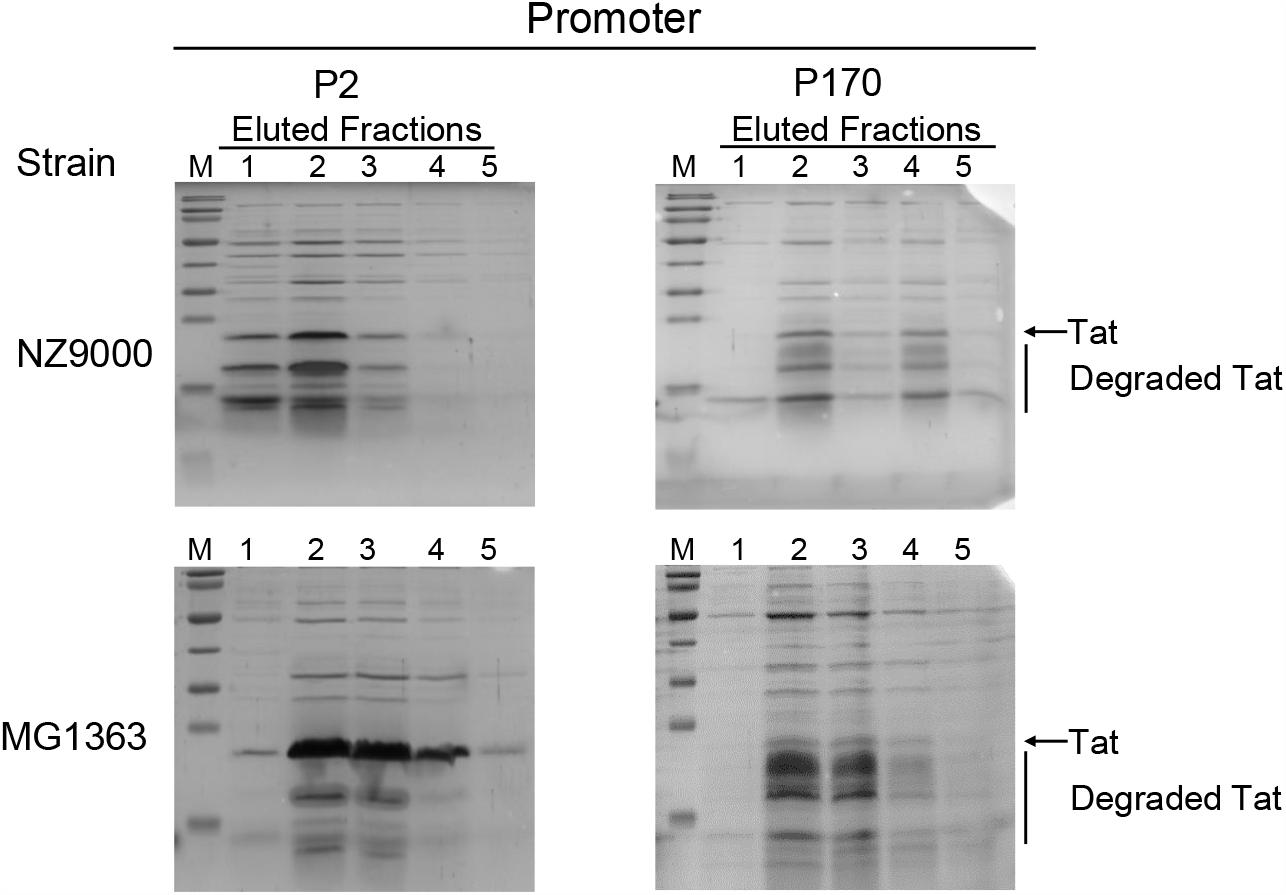
The expression profiles of Tat in two different bacterial strains. Tat was expressed from the P2 and P170 promoters in two *L. lactis* strains, MG1363 and NZ9000. Bacterial cultures of four litres were harvested at an OD of 1.4. Cells were treated with lysozyme at 37°C for one hour, followed by three cycles of sonication. The Tat protein was subsequently purified using Ni-NTA chromatography and eluted in five fractions of 500 μl each. Lane 1 represents marker (M), while Lanes 2, 3, 4, 5, and 6 represent the eluted fractions (30 μl each). The data are representative of at least three independent experiments.

### 3.3 The combined effect of Glucose concentration and pH of the medium on Tat expression

Qi et al. previously demonstrated a positive association between bacterial growth and increasing glucose concentration and pH maintenance of the medium [24]. To this end, we sequentially evaluated the effects of these two parameters on Tat expression. First, we examined the influence of glucose concentration on bacterial culture. The MG1363 strain was electroporated with the P2-Tat vector and cultured in M17 medium supplemented with 0, 0.25, 0.5, 1, or 2% glucose. Bacterial growth was monitored every two hours for 24 hours (Fig. 3A, left panel), and Tat expression was assessed by Western blot at three time points: 4, 10, and 24 hours (Fig. 3A, right panel). Notably, in the absence of glucose, bacterial growth reached a plateau at 0.5 OD within 6 hours and did not increase further. In contrast, the presence of glucose, even at the lowest concentration of 0.25%, allowed bacterial growth to continue to a higher density before plateauing at an approximate 1.4 OD (Fig. 3A, left panel). Sánchez et al. previously demonstrated that the accumulation of lactic acid in the medium could inhibit bacterial growth[25].

**Figure 3:**
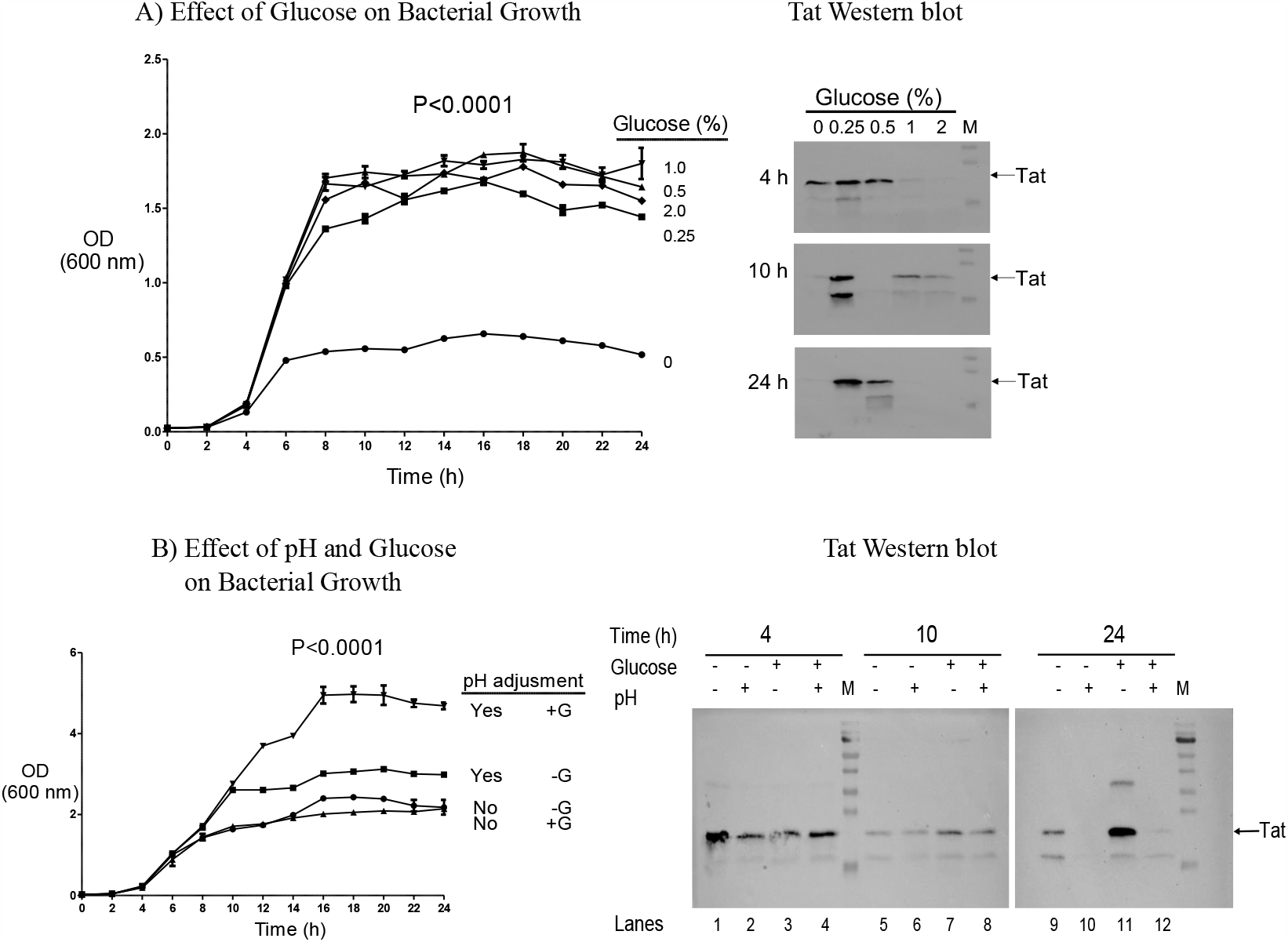
The effect of glucose and pH on Tat Expression in *L. lactis*. **(A)** Growth of *L. lactis* at varying concentrations of glucose. *L. lactis* cultures were grown at a glucose concentration ranging from 0% to 2%, and bacterial growth was monitored every two hours for 24 hours. Samples of 50 ml were harvested at 4, 10, and 24 hours for evaluating Tat expression by Western blot. **(B)** The combined influence of glucose concentration and pH on bacterial growth. The bacterial cells transfected with the Tat-expression vector were grown under four experimental conditions with or without glucose (+G and – G, respectively) and with or without adjusting the pH of the medium to 7.0 by adding 5 N NaOH every 2 hours. Cells were harvested, and Tat expression was monitored by Western blot. The data are representative of at least three independent experiments. Statistical analysis was performed using 2-way ANOVA and Bonferroni post-tests.

Western blot analysis of Tat at the 4 hours showed efficient protein expression at 0, 0.25, and 0.5% of glucose. Importantly, protein expression was significantly reduced at a higher concentration of glucose -for example, at 1% and 2% glucose concentrations, Tat expression was 11- and 53-folds lower, respectively, compared to 0.25% glucose (Fig. 3A, right panel). Notably, protein degradation was observed at all glucose concentrations by 10 hours, although Tat expression remained considerably high at 0.25% glucose. However, approximately half of Tat was degraded at 0.25% glucose. At 24 hours, Tat expression was observed mainly at 0.25% glucose, although a smaller quantity of protein expression was also seen at 0.5% glucose. In contrast, barely any Tat was expressed at other glucose concentrations. The Tat expression pattern reported here, including the protein degradation profile, is highly reproducible. As reported by others previously, Tat expression in L. lactis appeared to recapitulate a pulsed expression of recombinant proteins (see Discussion). In summary, we found efficient Tat expression at 0.25% glucose concentration at 4 and 24 hours, with minimal or no protein degradation. We next evaluated the effect of stabilizing the pH of the culture medium on bacterial growth and Tat expression at 0.25% glucose.

Lactococcal growth can progressively acidify the culture medium by secreting lactic acid,[24] therefore, adjusting the pH of the medium to a neutral pH is expected to enhance bacterial growth. In our hands, continuous bacterial culture reduced the pH of the medium to below 5.0 within six hours, by when, the optical density (OD) of the bacterial culture was stabilized at 1.4 and did not increase further even after 24 hours (Fig. 3A). To investigate the impact of pH adjustment on Tat expression, we cultured bacteria expressing Tat in the presence of 0.25% glucose, with or without a periodic pH adjustment of the culture medium. The pH of the medium was adjusted to 7.0 every 2 hours by adding 5 N NaOH solution. Additionally, after six hours, we also replenished the glucose concentration every 4 hours by adding 20% glucose stock solution. We maintained two additional bacterial cultures as controls, with or without adjusting the pH of the medium but without replenishing glucose in the medium.

The bacterial cells demonstrated comparable growth patterns under all experimental conditions up to 8 hours of culture. After this time, the adjustment of the pH of the medium to 7.0 appeared to have promoted bacterial growth more rapidly. For example, at 16 hours, the ODs of the bacterial cultures were 3.0 and 2.4 with or without pH adjustment, respectively (Fig. 3B, left panel). Furthermore, when the medium was supplemented with 0.25% glucose, a significantly higher level of bacterial growth was evident. Thus, when the medium pH was adjusted, at 16 hours, the OD of the bacterial cultures were 5.3 and 3.0 in the presence or absence of glucose, respectively. The data collectively suggested that the highest-level of bacterial growth could be seen in the presence of glucose and when the pH of the medium is adjusted to pH 7.0.

Subsequently, we monitored Tat expression using Western blot of the above bacterial cultures by harvesting the cells at three time points, 4, 10, and 24 hours. For the Western blot analysis, we could not estimate the total protein concentration of the cell lysates since the lysozyme used for cell lysis could interfere with the protein concentration estimation. Therefore, we used 0.4 OD equivalent of cell lysate in the Western blot analysis.

At 4 hours, neither glucose concentration nor pH adjustment appeared to have a significant effect on Tat expression, as under all four experimental conditions, comparable quantities of Tat expression were observed (Fig. 3B, right panel). At 10 hours, two- or three-folds higher levels of Tat expression were observed in glucose (compare lanes 5 and 6 with 7 and 8). Importantly, considerable differences in Tat expression profiles were evident at 24 hours. The highest-level Tat expression was observed in the presence of glucose but when the pH of the medium was not adjusted (Fig. 3B, right panel, lane 11). Surprisingly, 42-fold less Tat protein was expressed when the pH was adjusted, in the presence of glucose, although the bacterial growth was the highest under these conditions (Fig. 3B, right panel, compare lane 12 with 11). It is tempting to suggest that the activity of a bacterial protease active at the neutral pH could be responsible for intracellular Tat degradation (see Discussion). In summary, optimal Tat expression in *L. lactis* was seen in the presence of glucose and when the pH of the medium was acidic.

### 3.4 Intracellular Degradation of Tat

The previous data suggest that Tat is degraded by one or more cellular proteases, especially following prolonged culture. To confirm this observation, we cultured *L. lactis* harbouring the P2-Tat vector in the presence of 0, 0.25, 0.5, and 1 % glucose. The cultures were harvested at an early OD of 0.4 after 4 hours and at a late time point with an OD of 1.4 after 10 hours. Tat protein was purified from the harvested cultures and resolved in a 15% SDS-PAGE (Fig. 4A). Significantly, higher levels of Tat degradation were observed at the later points at all glucose concentrations. At 0.4 OD, relatively little or no Tat processing was observed, compared to OD 1.4, thus ascertaining our observations. Importantly, discrete Tat bands of low molecular weight, but not a protein smear, were evident in the gel, suggesting targeted processing of Tat by one or more specific cellular proteases.

**Figure 4:**
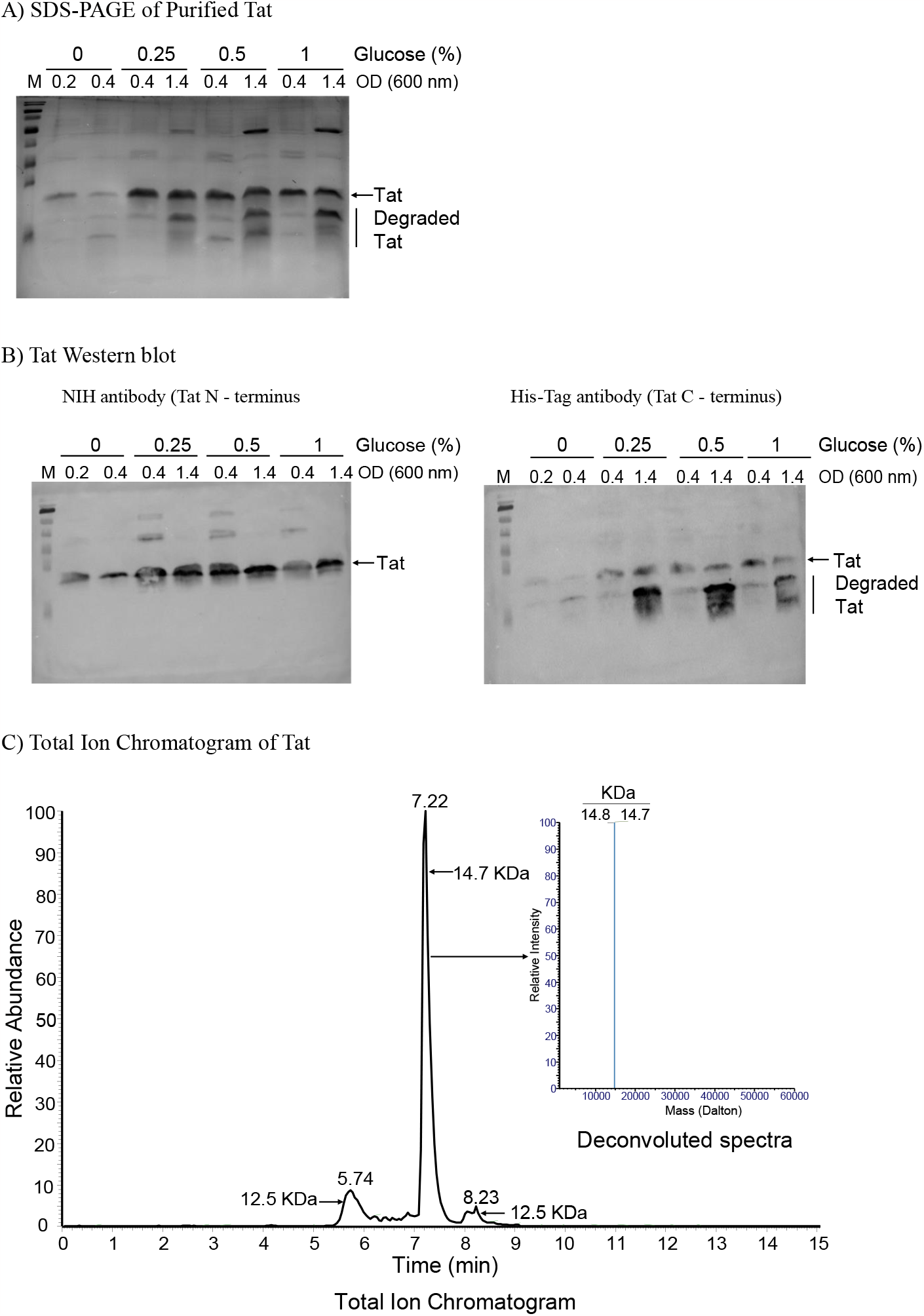
Analysis of Tat Degradation within cells. **(A)** SDS-PAGE analysis. *L. lactis* harbouring the P2-Tat vector was grown in a medium containing varying concentrations of glucose (0%, 0.25%, 0.5%, or 1%). Cultures were harvested at 0.4 or 1.4 OD, treated with lysozyme, and sonicated. Tat was purified using a Ni-NTA column, and 30 μl of the E2 fraction was subjected to SDS-PAGE and Western blot analyses. **(B)** Western blot analysis. The membrane was incubated first with an anti-Tat antibody 4138 (NIH AIDS Research and Reference Reagent Program) binding the N-terminal domain (left panel), stripped, and re-probed with a commercial antibody (Abcam, Cat. No. ab1178) binding the C-terminal His-Tag of Tat (right panel). **(C)** Mass spectrometry (MS) analysis of Tat expressed by bacteria grown in 0.25% glucose medium until reaching an optical density (OD) of 0.4. The expressed Tat protein was dialyzed with ultrapure water, and 200 ng of the protein was injected for analysis. The Total Ion Chromatogram (TIC) of Tat reveals three distinct peaks at different retention times: 5.74, 7.22, and 8.23 minutes. The peaks at 5.74 and 8.23 minutes correspond to the degraded products of Tat, while the peak at 7.22 minutes represents the intact mass of Tat. A deconvoluted image of the 7.22-minute peak is presented in the inset.

To understand the profile of Tat processing, we probed purified Tat in Western blot using two antibodies targeting two different epitopes in Tat. While the monoclonal antibody 4138 (NIH AIDS Research and Reference Reagent Program) binds the N-terminal domain of Tat, the His-Tag antibody (Abcam, Cat No. ab1178) binds the tag present at the C-terminal of Tat. The membrane was interrogated first with the anti-Tat antibody, stripped, and re-probed with the anti-His-Tag antibody. While the anti-Tat antibody detected a single band of 14.7 kDa of the entire Tat (Fig. 4B, left panel), the C-terminal binding anti-His-Tag antibody detected the full-length Tat as well as a processed band of smaller size (Fig. 4B, right panel). From this result, it is evident that Tat was subjected to intra-cellular processing after the N-terminal epitope.

We performed Mass-Spec analysis to assess the purity as well as the integrity of Tat. One microgram of purified Tat (under optimal conditions of 0.25% glucose and a harvested OD of 0.4) was dialysed with ultrapure water. The mass of the Tat protein was analysed using the deconvolution software (Biopharma Finder™ 3.2 Xtract algorithm), revealing three distinct retention times at 5.74, 7.22, and 8.23 minutes (Fig 4C). At 5.74 and 8.23 minutes, a fraction of degraded Tat protein (12.5 kDa) was observed, while at 7.22 minutes, a high intensity of intact 14.7 kDa protein was detected (Fig. 4C, inset). Mass-spec analysis confirmed the purification of Tat to the highest extent in a single step.

### 3.5 Tat expressed in *L. lactis* is competent for viral transactivation

Tat recruits the cellular positive transcriptional elongation factor (P-TEFb) to the viral transactivation response element (TAR)[26]. We evaluated the transactivation potential of the *L. lactis-*expressed Tat protein using two different cell models. First, we evaluated Tat in the HLM-1 cell line that harbours a Tat-defective HIV-1 provirus, which can be activated using functional Tat. One μg/ml of Tat protein was added to cells suspended in plain medium to activate the provirus. while 1X PBS served as the negative control. Four-hour post-transfection, the plain medium was replaced with a complete medium. Cultures were harvested after 48 hours, and the levels of p24 secreted into the medium were quantified using a commercial capture ELISA assay (J. Mithra Cat. No. IR232096). The secretion of p24 into the spent medium increased from 0.15 ng/ml to 1.726 ng/ml following Tat-treatment, a difference being statistically significant (Fig. 5A).

**Figure 5:**
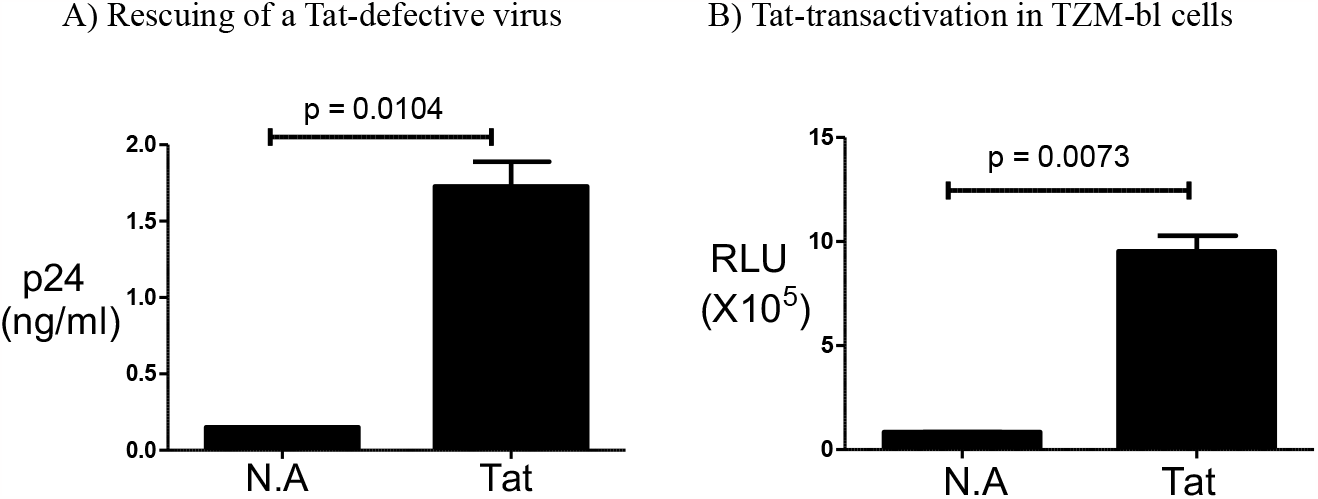
Functional validation of the *L. lactis* expressed Tat protein. **(A)** Rescue of a Tat-defective HIV-1 from the HLM-1 cells. HeLa-derived HLM-1 cells harbour an integrated full-length HXB2 virus containing an early stop codon within the Tat exon-1. In the presence of ectopic Tat, the virus is activated, releasing p24 into the medium. HLM-1 cells were treated with 1 μg/ml of the recombinant Tat protein. 1X PBS was used as the negative control. Following 48 hours of treatment, culture supernatants were collected to estimate p24 in ELISA. Statistical analysis was performed using Student’s paired t-test. The data are representative of at least three independent experiments. **(B)** Tat trans-activation in TZM-bl cells. TZM-bl cells harbour the β-Galactosidase and Luciferase Open Reading Frames (ORF) under the Long Terminal Repeat (LTR). In the presence of Tat protein, these cells express both luciferase and β-galactosidase. TZM-bl cells were treated with 1 μg/ml of recombinant Tat protein, 1X PBS was used as the negative control. After 24 hours of treatment, the cells were lysed with 1X passive lysis buffer, and the cell lysate was collected for Luciferase readings. Statistical analysis was performed using Student’s paired t-test. The data presented here is representative of at least three independent experiments.

The TZM-bl cell line, obtained from the NIH AIDS Research & Reference Reagent Program, is derived from HeLa cells that are CXCR4-positive. These cells have been further modified to express CD4 and CCR5 on their cell surface, and they contain integrated β-Galactosidase and a firefly luciferase reporter gene under the Long Terminal Repeat (LTR) elements. TZM-bl cells were transfected with one μg/ml of Tat protein using a commercial protein-transfection reagent (PierceTM Protein Transfection reagent, Cat. No. 89850), while 1XPBS was used for the negative control. After 24 hours of activation, the cells were lysed with 300 μl of 1X passive lysis buffer. Ten microlitres of cell lysate were mixed with 10 μl of Bright-Glo Luciferase substrate, and luciferase measurements were taken in a Luminometer. The luciferase value increased from 8.3X10^4^ to 9.5X10^5^ following the Tat treatment, the difference being statistically significant (Fig. 5B). Collectively, these data confirmed the transactivation competence of the recombinant Tat expressed in *L. lactis*. Apart from BL43 Tat, we also expressed YU2 Tat in *L. lactis* under similar conditions, and it was found to be functional (Data not shown).

## Discussion

The present study explored the potential of *Lactococcus lactis* as a host for the expression of Tat, a highly toxic protein of HIV-1, and optimized a few parameters crucial for efficient expression. In addition to governing viral transactivation, Tat controls several other functions, including modulating the host immune response, proliferation, apoptosis, angiogenesis, and others[27], [28]. The expression of substantial quantities of Tat is necessary to evaluate its various biological functions. However, the expression of Tat in a mammalian system presents a significant challenge given its inherently cytotoxic nature. Furthermore, Tat expression in *E. coli* also suffers from several technical limitations[10]. One major limitation of *E. coli* expression is the misfolding of the recombinant proteins, especially proteins such as Tat containing several cysteine residues. When we tested the lyophilized Tat produced in *E. coli* and *L. lactis* for their activity in HLM-1 cells, the activity of *E. coli*-expressed Tat was reduced by ninety percent compared to freshly isolated *E. coli* Tat protein (Data not shown). In the case of lyophilized Tat produced from *L. lactis*, it activated the HLM-1 cells as efficiently as freshly isolated Tat. Lactococcal expression of proteins, in contrast, may have a great advantage, especially when they contain multiple cysteine residues[26]. Incidentally, there are reports of successful expression of proteins containing multiple disulfide bonds in *L. lactis*[29] Another major limitation of *E. coli* expression of recombinant proteins is the danger of residual endotoxin contamination in the protein preparation. In the *E. coli* system, two rounds of purification are typically required to attain pure and functional Tat protein[10]. However, the process of endotoxin removal may lead to substantial loss of the recombinant protein.

To circumvent these challenges, we chose to express Tat in *Lactococcus lactis*. This approach is appealing since *L. lactis* is a gram-positive bacterium, and therefore the issue of endotoxin contamination is absent or minimal[27]. Another significant advantage of the *Lactococcus lactis* system is its ability to secrete proteins directly into the culture medium, thus simplifying the purification process.

The nisA promoter was shown to promote a robust and tightly controlled expression of recombinant proteins in *L. lactis*[18]. Since our efforts at expressing Tat under the nisA promoter were not successful, we evaluated other bacterial promoters. We could generate high-level Tat expression under the P2 promoter, which is a constitutive promoter. The pH-inducible P170 prompter also produced comparable quantities of Tat.

We faced the challenge of intracellular degradation of Tat under all experimental conditions, although Tat degradation was minimal when expressed in the MG1363 strain under the P2 promoter (Fig. 2). Several previous publications reported the problem of protein degradation in *L. lactis*. For example, Bera et al reported the degradation of streptokinase in *L. lactis* when expressed under the p170 promoter[14]. Bacterial culture leading to the stress of lactic acid accumulation upregulated HtrA, a surface protease, causing streptokinase degradation. Bera et al could resolve the degradation problem by utilizing an HtrA knockout strain. Likewise, Bermúdez-Humarán et al. observed the degradation of the cytoplasm-expressed Human Papillomavirus Type16 E7 protein in wildtype *L. lactis* but not in ΔClpP NZ9000 or ΔClpP MG1363 strains.

Several cellular proteases have been identified in *Lactococcus lactis*[30]. ClpP, an ATP-dependent enzyme, is the primary intracellular housekeeping protease in *L. lactis*[31]. In contrast, HtrA, a trypsin-like serine protease, is essential for bacterial growth at high temperatures. HtrA plays a role in degrading misfolded proteins at the cell surface[32]. Our results allude to the presence of a cytoplasmic protease, different from the above two, causing Tat processing, which must be identified and inactivated for efficient expression of recombinant proteins in this system. Several bacterial strains where such bacterial proteins have been inactivated to improve protein expression. For example, several HtrA knockout strains have been generated, such as the IL1403 HtrA mutant strain[33], the NZ9000 HtrA mutant strain[32], and the MG1363 HtrA mutant strain[11] . These knockout strains have proven effective in achieving the stable secretion of proteins into the culture medium[12].

Our attempts to enhance Tat yield by increasing bacterial OD proved counterintuitive as the prolonged bacterial culture resulted in augmented levels of Tat processing (Fig. 3B right panel). Probing with antibodies binding different epitopes of Tat, we show that Tat is processed after the N-terminal domain (Fig. 4B) by an unknown intracellular protease. In our experiments, we successfully produced functionally active Tat in the MG1363 strain under the P2 promoter in a medium supplemented with 0.25% glucose, and the bacterial cells were harvested after 4 hours at an OD of 0.4. However, when purified at 24 hours, we observed increased Tat degradation.

We explored additional options for enhancing Tat yield but without much success. We attached the USP45TM signal peptide 35 to Tat for secretion into the culture medium to prevent or minimize degradation. Following this manipulation, Tat was expressed at high concentration, up to 3 mg/l of protein/l in static flask cultures without degradation (data not shown). However, the expressed protein was not biologically active, probably due to rapid oxidation. Tat production under controlled fermentation conditions, minimizing oxidation, perhaps could be a potential solution. Over-expression of PmpA in *L. lactis* could also prove beneficial for enhancing protein stability. PmpA, a lipoprotein belonging to the peptidyl-prolyl cis/trans isomerase family[12] helped in the secretion of Staphylococcus hyicus lipase in *L. lactis* without experiencing degradation[34].

Our findings offer valuable insights into the expression of toxic proteins, such as HIV-1 Tat, in the static flask culture of *L. lactis*. The experimental conditions optimized here will serve as a useful guide for the cloning, expressing, and purifying difficult-to-express/toxic proteins in *L. lactis*. For proteins that are not prone to oxidation, like Tat, the addition of a signal peptide could make the protein secrete directly into the culture media.

## Conclusion

In summary, we successfully expressed the HIV-1 BL43 Tat protein using the P2-Tat-pNZ8148 plasmid in the MG1363 strain of *Lactococcus lactis*. When the culture was grown in the presence of 0.25% glucose and harvested at an OD of 0.4, we achieved an increased yield of intact Tat protein with minimal protein degradation. Although *L. lactis* produced approximately four-fold less Tat (0.5 mg/L) than *E. coli*, the purified protein lacked endotoxins. Unlike the *E. coli* expression system, which requires a dual purification process, Tat purification from *L. lactis* requires only a single step, resulting in significantly improved purity levels.

## Funding sourse

This work was supported by the Science and Engineering Research Board, Government of India (Sanction order no. CRG/2019/000820).

## Credit Author Contributions

DS designed, performed, interpreted, and wrote the manuscript. AD, YG, JS, and AP assisted in experimental design, while LH and UR were involved in experimental design, interpretation, and Manuscript corrections.

## Author statement

All authors acknowledge the following four statements to be true:

1. All authors concur with the submission.
2. The work has not been published elsewhere, either completely, in part, or in another form.
3. The manuscript has not been submitted to another journal and will not be published elsewhere Within one year after its publication in this journal.
4. The manuscript does not contain experiments using animals.

## Data statement

Raw data generated over the course of experiments is available upon reasonable request.

## AI statement

N/A

## Declaration of Interest Statement

The authors declare no conflict of interest.

## Data availability

Data will be made available on request.

## Acknowledgements

We express gratitude to ICMR for the fellowship and JNCASR for both intramural and extramural funding. Special thanks to Dr. Guhan Jayaraman for providing reagents and valuable suggestions.

## Notes

### Competing Interest Statement

The authors have declared no competing interest.

## References

1. Hemelaar J, Elangovan R, Yun J, Dickson-Tetteh L, Fleminger I, Kirtley S, Williams B, Gouws-Williams E, Ghys PD, Alash’le G A, Agwale S. Global and regional molecular epidemiology of HIV-1, 1990–2015: a systematic review, global survey, and trend analysis. The Lancet infectious diseases. 2019 Feb 1;19(2):143–55.

2. Spector C, Mele AR, Wigdahl B, Nonnemacher MR. Genetic variation and function of the HIV-1 Tat protein. Medical Microbiology and Immunology. 2019 Apr 1;208:131–69.

3. Pugliese A, Vidotto V, Beltramo T, Petrini S, Torre D. A review of HIV-1 Tat protein biological effects. Cell Biochemistry and Function: Cellular biochemistry and its modulation by active agents or disease. 2005 Jul;23(4):223–7.

4. Ben Haij N, Planès R, Leghmari K, Serrero M, Delobel P, Izopet J, BenMohamed L, Bahraoui E. HIV-1 Tat protein induces production of proinflammatory cytokines by human dendritic cells and monocytes/macrophages through engagement of TLR4-MD2-CD14 complex and activation of NF-κB pathway. PloS one. 2015 Jun 19;10(6):e0129425.

5. Ben Haij N, Leghmari K, Planès R, Thieblemont N, Bahraoui E. HIV-1 Tat protein binds to TLR4-MD2 and signals to induce TNF-α and IL-10. Retrovirology. 2013 Dec;10(1):1–2.

6. Abbas W, Herbein G. T-cell signaling in HIV-1 infection. The open virology journal. 2013;7:57.

7. Mueller N, Pasternak AO, Klaver B, Cornelissen M, Berkhout B, Das AT. The HIV-1 Tat protein enhances splicing at the major splice donor site. Journal of Virology. 2018 Jul 15;92(14):10–128.

8. El-Amine R, Germini D, Zakharova VV, Tsfasman T, Sheval EV, Louzada RA, Dupuy C, Bilhou-Nabera C, Hamade A, Najjar F, Oksenhendler E. HIV-1 Tat protein induces DNA damage in human peripheral blood B-lymphocytes via mitochondrial ROS production. Redox biology. 2018 May 1;15:97–108.

9. Clark E, Nava B, Caputi M. Tat is a multifunctional viral protein that modulates cellular gene expression and functions. Oncotarget. 2017 Apr 4;8(16):27569.

10. Siddappa NB, Venkatramanan M, Venkatesh P, Janki MV, Jayasuryan N, Desai A, Ravi V, Ranga U. Transactivation and signaling functions of Tat are not correlated: biological and immunological characterization of HIV-1 subtype-C Tat protein. Retrovirology. 2006 Dec;3(1):1–20.

11. Tucureanu MM, Rebleanu D, Constantinescu CA, Deleanu M, Voicu G, Butoi E, Calin M,MaduteanuI. Lipopolysaccharide-induced inflammation in monocytes/macrophages is blocked by liposomal delivery of Gi-protein inhibitor. International journal of nanomedicine. 2018 Dec 20:63–76.

12. Morello E, Bermudez-Humaran LG, Llull D, Sole V, Miraglio N, Langella P, Poquet I. Lactococcus lactis, an efficient cell factory for recombinant protein production and secretion. Microbial Physiology. 2007 Oct 24;14(1-3):48–58.

13. Frelet-Barrand A. Lactococcus lactis, an attractive cell factory for the expression of functional membrane proteins. Biomolecules. 2022 Jan 22;12(2):180.

14. Douillard FP, O’Connell-Motherway M, Cambillau C, van Sinderen D. Expanding the molecular toolbox for Lactococcus lactis: construction of an inducible thioredoxin gene fusion expression system. Microbial cell factories. 2011 Dec;10(1):1–0.

15. Bera S, Thillai K, Sriraman K, Jayaraman G. Process strategies for enhancing recombinant streptokinase production in Lactococcus lactis cultures using P170 expression system. Biochemical Engineering Journal. 2015 Jan 15;93:94–101.

16. Martinez-Jaramillo E, Garza-Morales R, Loera-Arias MJ, Saucedo-Cardenas O, Montes-de-Oca-Luna R, McNally LR, Gomez-Gutierrez JG. Development of Lactococcus lactis encoding fluorescent proteins, GFP, mCherry and iRFP regulated by the nisin-controlled gene expression system. Biotechnic & Histochemistry. 2017 Apr 3;92(3):167–74.

17. Jeeva P, Shanmuga Doss S, Sundaram V, Jayaraman G. Production of controlled molecular weight hyaluronic acid by glucostat strategy using recombinant Lactococcus lactis cultures. Applied microbiology and biotechnology. 2019 Jun 4;103:4363–75.

18. Eichenbaum Z, Federle MJ, Marra D, de Vos WM, Kuipers OP, Kleerebezem M, Scott JR. Use of the lactococcal nisA promoter to regulate gene expression in gram-positive bacteria: comparison of induction level and promoter strength. Applied and environmental microbiology. 1998 Aug 1;64(8):2763–9.

19. Jørgensen CM, Vrang A, Madsen SM. Recombinant protein expression in Lactococcus lactis using the P170 expression system. FEMS microbiology letters. 2014 Feb 1;351(2):170–8.

20. Zhu D, Liu F, Xu H, Bai Y, Zhang X, Saris PE, Qiao M. Isolation of strong constitutive promoters from Lactococcus lactis subsp. lactis N8. FEMS microbiology letters. 2015 Aug 1;362(16):fnv107.

21. Rud I, Jensen PR, Naterstad K, Axelsson L. A synthetic promoter library for constitutive gene expression in Lactobacillus plantarum. Microbiology. 2006 Apr;152(4):1011–9.

22. Wegkamp A, van Oorschot W, de Vos WM, Smid EJ. Characterization of the role of para-aminobenzoic acid biosynthesis in folate production by Lactococcus lactis. Applied and environmental microbiology. 2007 Apr 15;73(8):2673–81.

23. Sahoo TK, Jayaraman G. Co-culture of Lactobacillus delbrueckii and engineered Lactococcus lactis enhances stoichiometric yield of d-lactic acid from whey permeate. Applied microbiology and biotechnology. 2019 Jul 20;103:5653–62.

24. Qi W, Li XX, Guo YH, Bao YZ, Wang N, Luo XG, Yu CD, Zhang TC. Integrated metabonomic-proteomic analysis reveals the effect of glucose stress on metabolic adaptation of Lactococcus lactis ssp. lactis CICC23200. Journal of dairy science. 2020 Sep 1;103(9):7834–50.

25. Sánchez C, Neves AR, Cavalheiro J, dos Santos MM, García-Quintáns N, López P, Santos H. Contribution of citrate metabolism to the growth of Lactococcus lactis CRL264 at low pH. Applied and Environmental Microbiology. 2008 Feb 15;74(4):1136–44.

26. Tauer C, Heinl S, Egger E, Heiss S, Grabherr R. Tuning constitutive recombinant gene expression in Lactobacillus plantarum. Microbial cell factories. 2014 Dec;13(1):1–1.

27. Huigen MC, Kamp W, Nottet HS. Multiple effects of HIV-1 trans-activator protein on the pathogenesis of HIV-1 infection. European journal of clinical investigation. 2004 Jan;34(1):57–66.

28. Rubartelli A, Poggi A, Sitia R, Zocchi MR. HIV-1 Tat: a polypeptide for all seasons. Immunology today. 1998 Dec 1;19(12):543–5.

29. Singh SK, Tiendrebeogo RW, Chourasia BK, Kana IH, Singh S, Theisen M. Lactococcus lactis provides an efficient platform for production of disulfide-rich recombinant proteins from Plasmodium falciparum. Microbial cell factories. 2018 Dec;17:1–3.

30. Cortes-Perez NG, Poquet I, Oliveira M, Gratadoux JJ, Madsen SM, Miyoshi A, Corthier G, Azevedo V, Langella P, Bermúdez-Humarán LG. Construction and characterization of a Lactococcus lactis strain deficient in intracellular ClpP and extracellular HtrA proteases. Microbiology. 2006 Sep;152(9):2611–8.

31. Frees D, Ingmer H. ClpP participates in the degradation of misfolded protein in Lactococcus lactis. Molecular microbiology. 1999 Jan;31(1):79–87.

32. Bermudez-Humaran LG, Langella P, Miyoshi A, Gruss A, Guerra RT, Montes de Oca-Luna R, Le Loir Y. Production of human papillomavirus type 16 E7 protein in Lactococcus lactis. Applied and environmental microbiology. 2002 Feb;68(2):917–22.

33. Poquet I, Saint V, Seznec E, Simoes N, Bolotin A, Gruss A. HtrA is the unique surface housekeeping protease in Lactococcus lactis and is required for natural protein processing. Molecular microbiology. 2000 Mar;35(5):1042–51.

34. Drouault S, Anba J, Bonneau S, Bolotin A, Ehrlich SD, Renault P. The peptidyl-prolyl isomerase motif is lacking in PmpA, the PrsA-like protein involved in the secretion machinery of Lactococcus lactis. Applied and environmental microbiology. 2002 Aug;68(8):3932–42.

